# Deep plasma proteomics with data-independent acquisition: A fastlane towards biomarkers identification

**DOI:** 10.1101/2024.02.23.581160

**Authors:** Bradley Ward, Sébastien Pyr dit Ruys, Jean-Luc Balligand, Leïla Belkhir, Patrice D. Cani, Jean-François Collet, Julien De Greef, Joseph P. Dewulf, Laurent Gatto, Vincent Haufroid, Sébastien Jodogne, Benoît Kabamba, Maxime Lingurski, Jean Cyr Yombi, Didier Vertommen, Laure Elens

## Abstract

Plasma proteomic is a precious tool in human disease research, but requires extensive sample preparation in order to perform in-depth analysis and biomarker discovery using traditional Data-Dependent Acquisition (DDA). Here, we highlight the efficacy of combining moderate plasma prefractionation and Data-Independent Acquisition (DIA) to significantly improve proteome coverage and depth, while remaining cost- and time-efficient.

Using human plasma collected from a 20-patient COVID-19 cohort, our method utilises commonly available solutions for depletion, sample preparation, and fractionation, followed by 3 LC-MS/MS injections for a 360-minutes DIA run time. DIA-NN software was then used for precursor identification, and the QFeatures R package was used for protein aggregation.

We detect 1,321 proteins on average per patient, and 2,031 unique proteins across the cohort. Filtering precursors present in under 25% of patients, we still detect 1,230 average proteins and 1,590 unique proteins, indicating robust protein identification. Differential analysis further demonstrates the applicability of this method for plasma proteomic research and clinical biomarker identification.

In summary, this study introduces a streamlined, cost- and time-effective approach to deep plasma proteome analysis, expanding its utility beyond classical research environments and enabling larger-scale multi-omics investigations in clinical settings.

## Introduction

### The human plasma proteome

Human plasma proteomics is a rapidly evolving area of research aiming to identify and understand the functions of proteins present in human blood, which virtually gives a holistic view of the human body and its health status. Containing a wide range of substances, including hormones, metabolites, and proteins produced by numerous organs and tissues within the body, it provides a snapshot of a person’s overall health, as many diseases can have systemic effects on the body that are reflected in blood ^1^. There are currently 20,389 predicted coding genes, with strong protein-level evidence for 18,397 proteins in the human proteome ^2^. Of these, just over 4,500 canonical proteins have been detected in human plasma, meaning this biofluid represents around 20% of the human proteome currently detectable ^3^. Due to a combination of its ease of collection and biological significance, plasma is highly advantageous to disease research, for example, much of the research into the effects of coronavirus disease 2019 (COVID-19) has been conducted through plasma proteomic studies which have been determinant in identifying biomarkers such as levels of S100A9 ^4,5^, ITIH2 ^6^, TNF-α ^7^, and cytokines ^8^, elucidating disease pathophysiology by discovering dysregulated proteins and pathways, and finding potential drug targets and treatments.

As useful as it is to study the human plasma proteome, it is also a challenging analytical task for several reasons. Firstly, plasma is an incredibly heterogeneous sample, containing lipids, metabolites, and other small molecules which must be removed to prevent interference with protein detection and quantification. Secondly, its protein content is incredibly complex, inter-individually variable, and dynamic, with proteins concentrations spanning 12-13 orders of magnitude ^9^. This is outside the dynamic range of modern analytical instruments, making detection and quantification of low-abundant proteins difficult ^10^. Thirdly, many proteins undergo post-translational modifications which generate a myriad of proteoforms requiring specialized techniques and equipment to identify. And lastly, analysing the large datasets generated by these studies requires sophisticated bioinformatics and biostatistical tools as well as expertise to interpret and validate the data, requiring a multidisciplinary approach. Despite these challenges, advances in technology and methodology are providing new opportunities to explore the plasma proteome, identifying and quantifying an increasingly complex array of proteins, and facilitating a deeper understanding of disease mechanisms and potential therapeutic targets.

### Depletion workflows

Plasma proteins span a huge dynamic range, with 22 proteins accounting for 99% of the proteome by abundance; the remaining 1% of proteins are usually those of interest to researchers and clinicians. Depleting high abundant proteins can enhance the detection of low-abundance ones, but many low-abundant proteins may also be co-depleted as many high-abundant proteins have carrier functions ^11,12^. Albumin, for instance, binds to many peptides and proteins collectively referred to as the albuminome, comprising almost 250 direct and indirect interaction proteins ^13^. Other highly abundant proteins usually depleted include immunoglobulins, fibrinogen, and apolipoprotein, each with their own passenger proteins; collectively, the proteins comprising the depleted fraction have been termed the “depletome”^14^. Despite this off-target protein depletion, depletion is a critical strategy for any proteomic study which wishes to investigate deep into the human proteome. Additionally, as depletion does not completely remove proteins, high sensitivity techniques (such as DIA) may still be able to identify them, rendering this less of an issue.

### Fractionation workflows

Fractionation strategies, which can be used alone or in combination with depletion strategies, divide samples into fractions of lower dynamic range and complexity, helping to better identify low-abundant proteins. Fractionation can take place before and/or after digestion, at the protein and/or peptide level, although is most commonly performed at the peptide level for bottom-up proteomics; the fractionation discussed in this paper will refer to peptide-level fractionation. Generally, more fractions result in more proteins identified, with one study showing increases of 90%, 48%, and 32% unique proteins identified at 5 vs 20, 12 vs 24, and 20 vs 40 fractions respectively ^15^. However, drawbacks include minor sample loss and increased analytical time, which can greatly increase costs.

Multiple approaches have been utilised, from strong cation exchange chromatography to high pH reverse-phase chromatography approach ^16^. These methods are most commonly executed “off-line”, whereby samples are first fractionated, followed by further separation via liquid chromatography prior to mass spectrometry.

### Analytical technologies

Plasma proteomic analysis employs various methods, with mass spectrometry being the most common. The high sensitivity and specificity of this technique is capable of detecting and identifying hundreds to thousands of proteins in a single analysis ^9^.

Traditionally, untargeted mass spectrometry proteomics relied on data-dependent acquisition (DDA), but its selectivity and reproducibility have limitations due to its stochastic nature, and so, DIA techniques have grown in popularity (the differences of the two techniques can be seen in figure 1). Many DIA techniques have been introduced ^17–21^, but the general results are that all precursor product ions are analysed in a systematic and unbiased manner, allowing DIA to achieve reproducibility rates of up to 99% and 98% at the peptide and protein level respectively, compared to around 80% for DDA methods ^22^. However, DIA usually results in highly complex MS2 spectra which are challenging to interpret and quantify, requiring specialised software and computational resources. Much software has been developed to aid in DIA bioinformatic work in recent years, the most popular being Skyline ^23^, OpenSWATH ^24^, Spectronaut ^25^, DIA-Umpire ^26^, and DIA-NN ^27^.

**Figure 1.**
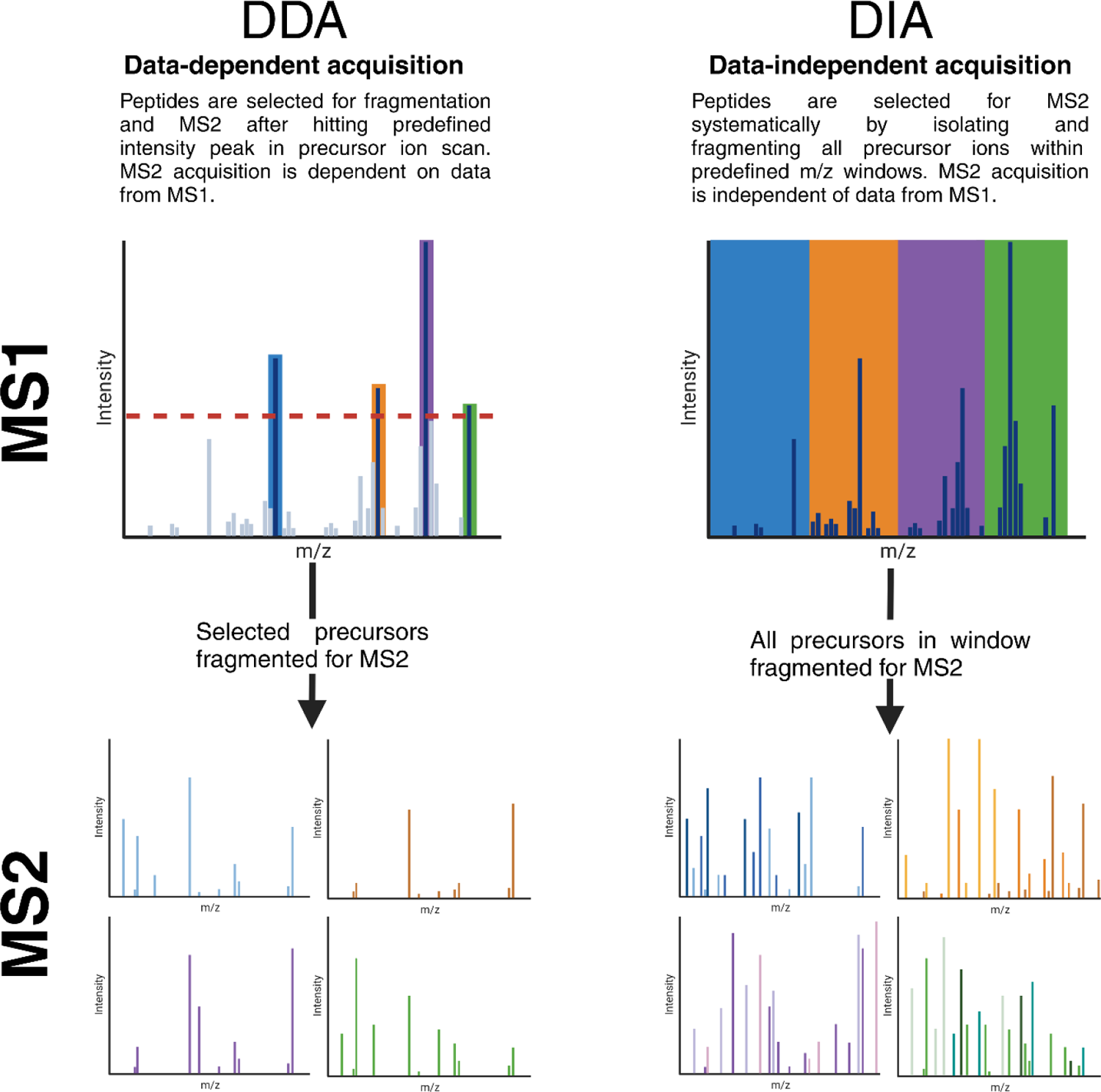
Difference between DDA and DIA. In DDA, precursors are selected for fragmentation and MS2 acquisition based on the data in the MS1 precursors ion scan. In this case, only the selected peptides are fragmented, leading to simpler MS2 spectra and easier data analysis. In DIA, precursor fragmentation selection is independent of MS1 data, instead, all precursors within a predefined m/z window/range are fragmented together for MS2 acquisition. This leads to more complex MS2 spectra in which multiple precursors are present and so more powerful bioinformatic tools must be used to identify individual precursors. Created with BioRender.com.

Specifically, for DIA-NN, it employs a peptide-centric method which is particularly suitable for DIA experiments, as compared to more traditional spectrum-specific methods. Ting (. (2015), have written a comprehensive review outlining the differences between these two approaches in the context of DIA.

Despite the advantages of DIA, it is not currently the first method of choice if looking to identify the maximum number of proteins from a small collection of complex samples, as the proteomic depth of DDA-based methods can be significantly increased when coupled with extensive fractionation and long-gradient liquid chromatography (LC). Regardless of which DDA or DIA strategies are used, the quality of the data received heavily depends on the sample preparation employed prior to LC-MS injection.

### Summary

Considering all the steps involved in a proteomic study, costs per sample can quickly increase. Here we have briefly discussed protein depletion and fractionation strategies, but other techniques such as isobaric labelling can also be involved in order to allow for multiplexing and increased sample throughput. As most studies do not share per sample costs, it can be difficult to get an accurate price regarding the different techniques used and data quality, but generally, as sample preparation increases in complexity, price and data quality usually increase in parallel. Although well known to academics, the literature is lacking significantly in discussions surrounding pricing per patient/sample for various experimental techniques. Optimising funding can be one of the most significant ways in which research itself can be optimised, allowing for more in-depth experiments or increasing cohorts limited by funding size. Here we present a simplified sample preparation protocol consisting of depletion followed by fractionation and subsequent pooling of fractions, resulting in 3 LC-MS/MS injections per sample. Coupled with the recent developments in DIA-based proteomics and DIA analytical software, we are able to achieve an average of 1,231 proteins identified per sample and 1,603 unique proteins in a 20-patient preliminary cohort (described below), at a much lower cost-per-sample than highly fractionated DDA methodologies, using widely available commercial kits and solutions.

## Methods

### Proof-of-concept sample cohort

The cohort included in this study comprises a 20 patient sub-cohort from the HYGIEIA study^29^. Patients were enrolled from August 2020 to December 2023. The protocol has received ethical approbation from the local ethical committee (2021/30DEC/543) and has been registered on clinicaltrial.gov (NCT05557539). During recruitment, a subset of 20 patients were selected for a validation cohort and split into five balanced groups (n=4): healthy controls, respiratory infection controls, moderate COVID-19 WHO score <=2, severe COVID-19 WHO score 3-5, and critical COVID-19 WHO score >=6 ^30^. Each group was comprised of two males and two females, average age of the cohort was 47 years old at hospitalisation, average BMI was 26kg/m^2^. 16 patients were of Caucasian ethnicity, with the remaining four patients having North African, Caribbean, Arab, and unknown ethnicities (self-reported by patient).

### Patient sample storage

Patient samples were collected during the acute phase of the infection and/or within one day of patient consent to study inclusion. Plasma samples were collected in 4mL Lithium heparin gel prepared S-Monovette tubes (Sarstedt, Nümbrecht, Germany) and kept at 4°C for a maximum of 4 hours until plasma was separated via centrifugation (2600g for 10 minutes at 4°C) (Sigma 3-16KL, Sigma Laboratory Centrifuges, Osterode am Harz, Germany), then aliquoted, snap frozen in liquid nitrogen, and stored at −80°C until later processing.

### Protein and peptide quantitation

In the following methodology, where appropriate, protein quantitation was performed using the Bio-Rad Protein assay kit II (Bio-Rad Laboratories, Hercules, United State) and peptide quantitation was performed using the Pierce™ Quantitative Peptide Assays & Standards (Thermo Fisher Scientific, Waltham, United States).

### Protein depletion

Protein depletion was performed according to manufacturer instructions. Briefly, the High Select™ Top14 Abundant Protein Depletion Resin (Thermo Fisher Scientific, Waltham, United States) is gently homogenised by inverting the tube several times. Then 600µl of resin is deposed into a 2mL Protein LoBind® Eppendorf tube (Eppendorf, Hamburg, Germany) and allowed to equilibrate to room temperature (RT) (15°C – 25°C). 22µl of plasma sample are then added and gently mixed via inversion before adding the tube on a rotator (Loopster digital, IKA, Staufen, Germany) at 12 RPM for 10 minutes at room temperature (RT, 18-25°C).

The plasma-resin mixture is then transferred to a Pierce™ Micro-Spin Column (Thermo Fisher Scientific, Waltham, United States) placed in a 2mL Protein LoBind® Eppendorf tube and centrifuged at 1000g for 2 minutes at RT (Microcentrifuge 5415, Eppendorf, Hamburg, Germany).

### Digestion

The flow through is first incubated at 95°C for 5 minutes (ThermoMixer C, Eppendorf, Hamburg, Germany), then the sample is allowed to cool for 5 minutes at RT. 33µl of 50mM DL-Dithiothreitol (5mM final concentration) (Sigma-Aldrich, St. Louis, United States) are then added to the sample and incubated at 56°C for 1 hour at 1000 RPM (ThermoMixer C, Eppendorf, Hamburg, Germany). 37µl of 500mM chloroacetamide (50mM final concentration) (Sigma-Aldrich, St. Louis, United States) are added and the mixture is vortexed briefly (max power, 3 seconds, Vortex-Genie 2, Scientific Industries, Bohemia, United States) and spun down (3 seconds, Mini Centrifuge, ExtraGene, Davis, United States), then incubated at RT for 30 minutes in the dark. Then 65.5µl 100% trichloroacetic acid (15% final concentration) (Sigma-Aldrich, St. Louis, United States) are added, the sample is then vortexed briefly and, again, spun down, and finally, incubated on ice for 30 minutes.

After incubation, the sample is centrifuged for 4000g for 7.5 minutes, then washed 3 times by removing supernatant, adding 500µl −20°C acetone (Carl Roth, Karlsruhe, Germany) sonicating at 37kHz pulsed for 2 minutes (Elmasonic P, Elma, Singen am Hoentwiel, Germany), then centrifuging at 4000g for 5 minutes. After repeating 3 times, the supernatant is again removed, and the sample is allowed to air-dry for 10 minutes at RT. Next, 75µl triethylammonium bicarbonate (TEAB) 50mM (Thermo Fisher Scientific, Waltham, United States) are added to the tube, followed by two rounds of sonication at 37kHz pulsed for 2 minutes each. The sample is subsequently vortexed and spun-down, then 2.5µg sequencing grade modified trypsin (V5117, Promega, Madison, United States) are added (50:1 protein:protease) and the sample is incubated overnight at 37°C at 750 RPM (ThermoMixer C).

### Fractionation

Following incubation, 8.3µl 1% trifluoroacetic acid (Biosolve, Dieuze, France) are added to the sample (0.1% final concentration), then 80µg total peptide are directly deposed on a Pierce™ High pH Reversed-Phase Peptide Fractionation column (Thermo Fisher Scientific, Waltham, United States) and centrifuged at 3000g for 2 minutes. The flow through is then discarded, and the column washed with 300µl MS grade water at 3000g for 2 minutes. Fractions are collected from column in 1.5mL Protein LoBind Eppendorf tubes (Eppendorf, Hamburg, Germany) by adding 300µl of the appropriate elution solution to the column and centrifuging at 3000g for 2 minutes. Elution solutions contain 0.1% triethylamine with either 7.5%, 10%, 12.5%, 15%, 17.5%, 20%, or 50% acetonitrile for collecting fractions 1 through 7 respectively. Following fractionation, pooling of the fractions are carried out with 100µl each of fraction 1, 2, and 5 as pooled fraction 1, 150µl each of fraction 3 and 6 as pooled fraction 2, and 150µl each of fraction 4 and 7 as pooled fraction 3. Pooling fractions in such a way helps ensure chemically dissimilar peptides are pooled together. The pooled fractions are then incubated on dry ice for 5 minutes, then open tubes are placed into a vacuum concentrator (SpeedVac SRF110, Thermo Fisher Scientific, Waltham, United States) at 4°C until pooled fractions have fully evaporated. 20µL MS resuspension buffer (3.5% ACN, 0.1% TFA) are subsequently used to resuspend each tube, and 3 sonication cycles at 37kHz pulsed for 2 minutes are repeated. Finally, each resuspended sample is briefly vortexed and spun down.

### DIA LC-MS/MS injection

1 µg of peptides dissolved in solvent A ((MS resuspension buffer) 0.1% TFA in 3.5% ACN) was directly loaded onto a reversed-phase pre-column (Acclaim PepMap 100, Thermo Fisher Scientific, Waltham, United States) and eluted in backflush mode. Peptide separation was achieved using a reversed-phase analytical EasySpray column (Acclaim PepMap RSLC C18, 0.075mm × 250mm, Thermo Fisher Scientific, Waltham, United States) with a 120 minute linear gradient of 4%–32% solvent B (0.1% TFA in 80% ACN) for 100 minutes, 32%–50% solvent B for 5 minutes, 50%–90% for 5 minutes and holding at 95% for the last 9 minutes at a constant flow rate of 300nL/min on an Ultimate 3000 RSLC nanoHPLC system (Thermo Fisher Scientific, Waltham, United States). The peptides were analyzed by an Orbitrap Exploris 240 mass spectrometer (Thermo Fisher Scientific, Waltham, United States) with enabled advanced peak determination (APD). The peptides were subjected to an EasySpray ionization source, followed by MS/MS in the Exploris 240 coupled online to the HPLC. Intact peptides were detected in the Orbitrap at a full width at half-maximum resolution of 30,000 (FWHM) and MS/MS spectra were acquired in the Orbitrap after stepped higher energy collisional dissociation (HCD) fragmentation at 22%, 26% and 30% of maximum. A data-independent procedure of MS/MS scans was applied for the precursor ions within a m/z range of 500-740 and an m/z isolation window of 4 at a resolution of 60,000 (FWHM). MS1 spectra were obtained with an automatic gain control (AGC) target of 1.2 × 10^6^ ions and a maximum injection time of 55ms. MS2 spectra were acquired with an AGC target of 1.5 × 10^5^ ions and a maximum injection time set to auto. For MS2 scans, the m/z scan range was set at 145 to 1450.

### DDA LC-MS/MS injection

Peptides were injected and separated as described above. The peptides were analyzed by an Exploris240 Orbitrap mass spectrometer (Thermo Fisher Scientific, Waltham, United States). The peptides were subjected to nanospray ion source followed by MS/MS in Exploris240 coupled online to the nano-LC. Intact peptides were detected in the Orbitrap at an FWHM resolution of 60,000. Peptides were selected for MS/MS using HCD setting at 30%; ion fragments were detected in the Orbitrap at an FWHM resolution of 15,000. A data-dependent procedure that alternated between one MS scan followed by MS/MS scans was applied for 3 seconds for ions above a threshold ion count of 1.0e^4^ in the MS survey scan with 40.0 seconds dynamic exclusion. The electrospray voltage applied was 2.1kV. MS1 spectra were obtained with an AGC target of 4e^5^ ions and a maximum injection time of 50ms, MS2 spectra were acquired with an AGC target of 5e^4^ ions and a maximum injection time set to dynamic. For MS scans, the m/z scan range was 375 to 1800.

### Bioinformatic analysis

DIA-NN was first used to generate an in-silico predicated spectral library using all peptides from the Human PeptideAtlas (2023-01 build) ^31^. Following HPLC-MS/MS acquisition, Thermo .raw spectra files for all fractions and all patients were imported into DIA-NN ^27^ using the Thermo MS File Reader (Thermo Fisher Scientific, Waltham, United States). The in-silico library was used as the spectral library, and to further aid peak detection, match-between-runs mode was enabled. In addition to default options, the following options were used: FASTA digest for library-free search/library generation; Deep learning-based spectra, RTs and IMs prediction; Maximum number of variable modifications = 1, Modifications = N-term M excision, C carbamidomethylation, Ox(M); Unrelated runs; MBR; Neural network classifier = Double-pass mode; Quantification strategy = Any LC (high accuracy); Precursor false discovery rate (%) = 1.0. After processing, the corresponding report file was then loaded into R for processing (v4.3.1) ^32^. The complete script can be found in Appendix 1, but in brief, runs were separated into pooled fraction 1, pooled fraction 2, and pooled fraction 3 to be processed separately via QFeatures(v3.17) ^33^, log transformed, and normalised. The 3 fraction assays were then combined and aggregated into protein groups via the robust summary methodology ^34^ The process for non-fractionated samples was similar, except that runs did not have to be separated and then recombined.

For DDA samples, the resulting MS/MS data was processed using Sequest HT search engine within Proteome Discoverer 2.5 SP1 (Thermo Fisher Scientific, Waltham, United States) against a Homo sapiens protein database obtained from Uniprot (32,314 entries), trypsin was specified as cleavage enzyme allowing up to 2 missed cleavages, 2 modifications per peptide and up to 5 charges. Mass error was set to 10 parts per million (ppm) for precursor ions and 0.02 Da for fragment ions. Oxidation on Met (+15.995 Da), pyro-Glu formation from Gln or Glu (−17.027 Da or −18.011 Da respectively) were considered as variable modifications. False discovery rate (FDR) was assessed using Percolator and thresholds for protein, peptide and modification site were specified at 1%. Label free quantification was performed by Proteome Discoverer 2.5 using the area under the curve (AUC) for each peptide.

During data analysis, predicted protein concentrations in plasma were obtained from the Plasma Proteome Database which reports plasma protein levels in healthy conditions as reported in literature ^35,36^. For COVID-19 biomarkers and COVID-19 associated proteins, the OncoMX COVID-19 biomarker database and OpenTargets platform (targets associated with COVID-19) were used respectively ^37,38^.

### Statistical analysis

Statistical analysis of the data was carried out using R(v4.3.1) ^32^. Two sample proportion tests and t test were used to evaluate the significance of the data when appropriate, such as when comparing the differences in the mean number of identified proteins in fractionated and non-fractionated samples. The types of test used are referenced within the text when needed alongside the adjusted p-values. For t tests comparing DDA to DIA, or non-fractionated to fractionated, the assumption is that DIA, or fractionation, produces superior results and so one-sided t tests are used in these cases. Significance levels of protein abundances was calculated using t tests. All code used to generate statistics can be found in appendix 1.

## Results

### DIA fractionation vs non-fractionation

Mean number of proteins detected and quantified in human plasma are found to be significantly higher, with an increase of around 50% for fractionated samples compared to non-fractionated samples (figure 2), with an average of 1,321 and 894 proteins per patients for fractionated and unfractionated samples respectively (right-tailed, paired, t-test, p-value = 1.17e-06). A similar trend is seen for total unique quantified proteins, with 2,031 total unique proteins found in fractionated samples and 1,765 in non-fractionated samples. The numbers of proteins detected in all patients was also significantly increased in fractionated samples, with 570 common proteins (around 43% of proteins on average) compared to 303 common proteins in non-fractionated samples (around 34% of proteins on average) (two sample proportion test, p-value = 1.38e-63).

**Figure 2.**
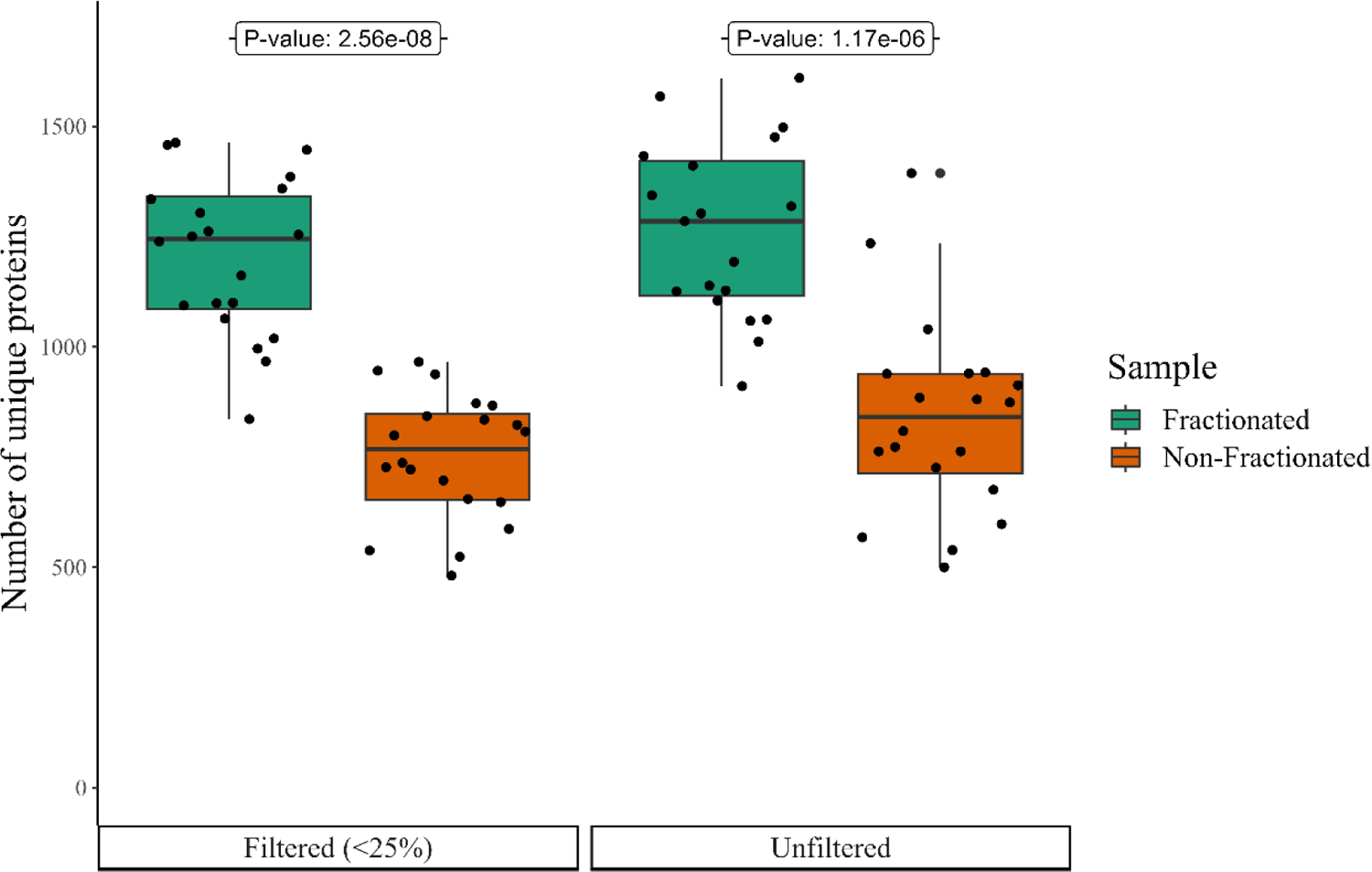
Number of proteins detected per patient in fractionated and non-fractionated samples. Black points represent individual patients, while boxplots represent the whole cohort. Right-tailed, paired, t-test. Filtered: t = 8.65, p-value = 2.56e-08. Unfiltered: t = 6.65, p-value = 1.17e-06. Filter removes proteins if present in less than 25% of patients

Filtering the data to investigate those proteins consistently found in the samples (by filtering precursors present in less than 25% of patients), results are similarly significant, with an average of 1,231 vs 788 detected and quantified proteins per patient respectively (right-tailed, paired, t-test, p-value = 2.56e-08). 1,590 total unique proteins found in fractionated samples and 1,063 in non-fractionated samples. And finally, there is only a small decrease in total common proteins (but a small relative increase compared to total unique proteins), with 565 vs 297 in fractionated and non-fractionated samples respectively (46% vs 38%) (two sample proportion test, p-value = 4.62e-46). Alongside the enhanced detection of proteins within fractionated samples, fractionation also appears to improve protein detection consistency between samples. This is visualised in figure 3, which shows a higher consistency of protein identification across patient samples with fractionation as opposed to non-fractionated samples.

**Figure 3.**
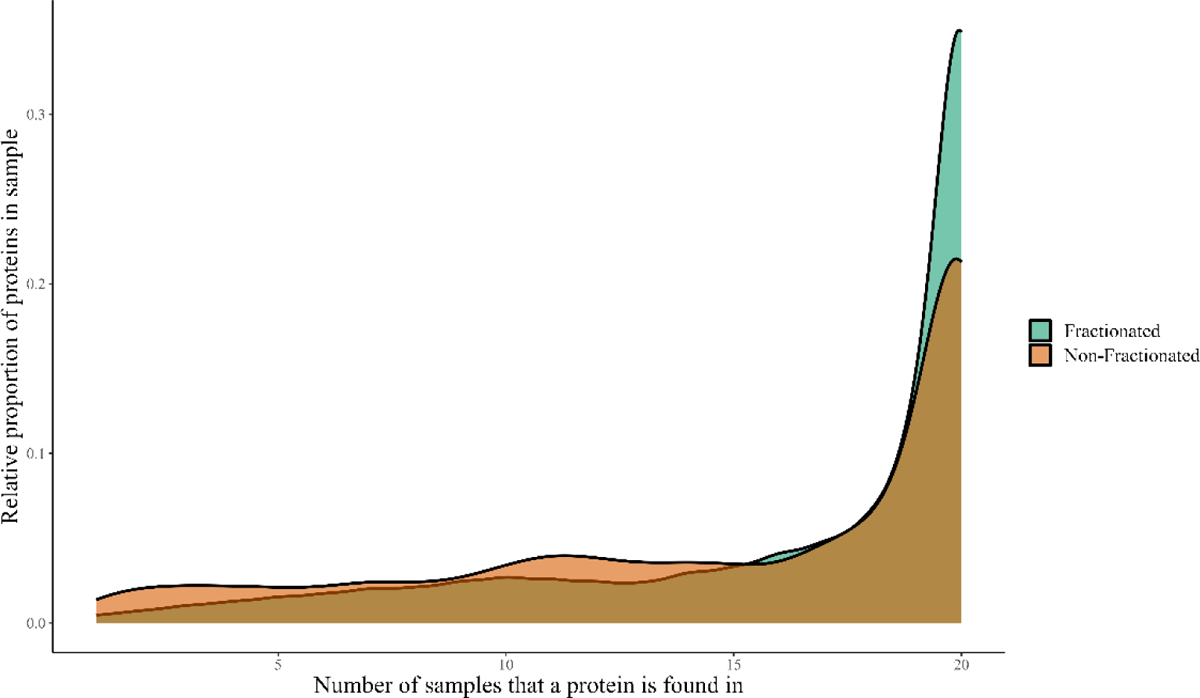
Density plot. Illustrates the proportion to which proteins are found in each sample within the cohort (20 patients) for each group. A higher peak towards the right of the graph suggests proteins are found more consistently in a higher proportion of the patient cohort.

Not only are more unique proteins found within fractionated samples, but a higher proportion of these proteins are found within a higher number of samples. Looking at figure 4, we can also see the majority of information (in terms of detected proteins) is almost completely found within the fractionated samples, with non-fractionated samples providing only a small number of unique proteins. Indeed, out of 2,061 unique proteins in both fractionated and non-fractionated samples, 296 are uniquely found in fractionated samples, 1,735 are found in both, and only 30 are found uniquely in non-fractionated samples. In terms of precursors, 1,864 are uniquely found in fractionated samples, 9,373 are commonly found, and 232 are uniquely found in non-fractionated samples. Looking at both fractionated and non-fractionated samples, a total of 11,469 distinct precursors were detected, 11,237 of these were found in fractionated samples, and 9,605 of these were found in the non-fractionated samples.

**Figure 4.**
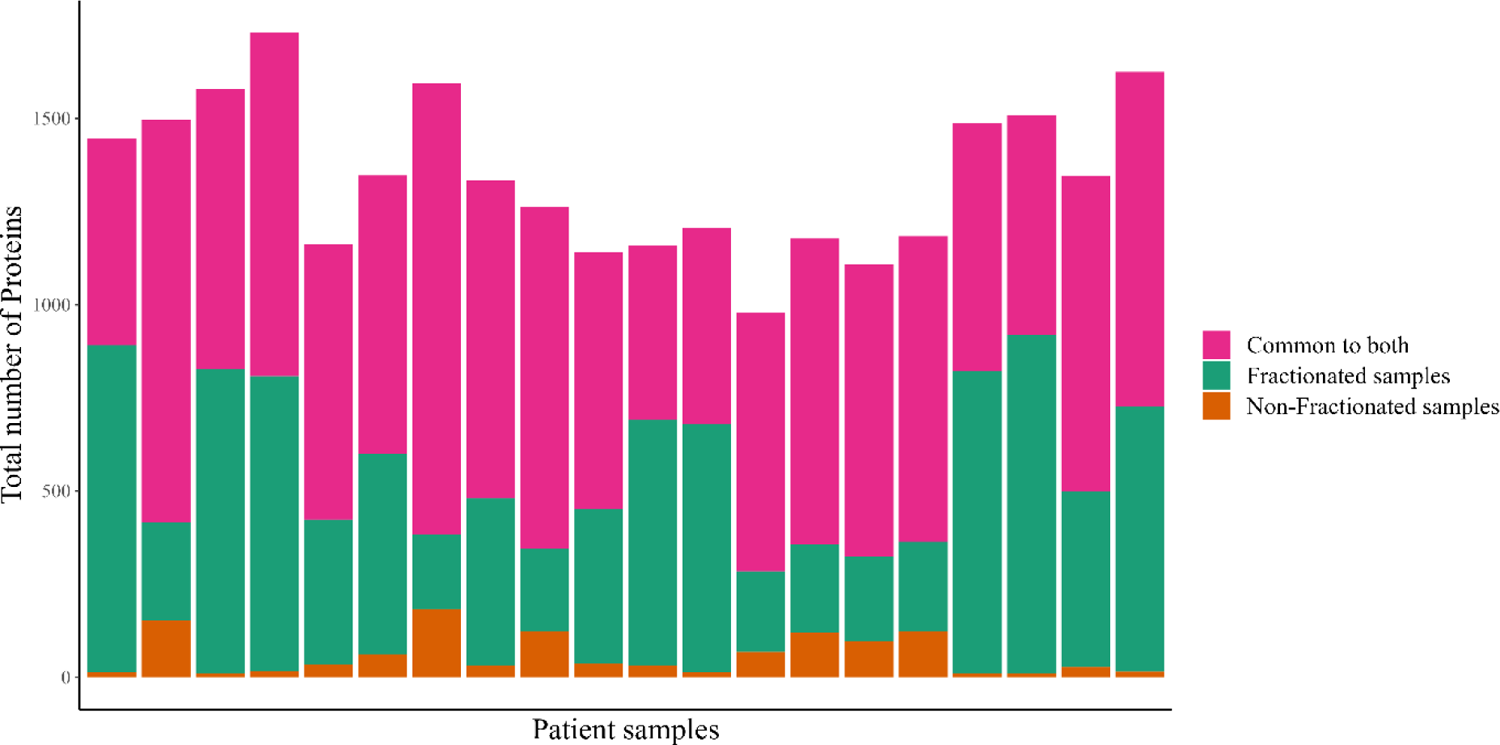
Bar plot. Number of proteins detected per patient, colour grouping within each bar shows which proportion of the proteins are found in both fractionated and non-fractionated samples (common to both) (pink), and which are unique to fractionated (green) or non-fractionated samples (orange).

After filtering as described above, in which precursors are removed if not found in more than 25% of samples, these numbers dropped to 8,841 total unique precursors, 8,627 in fractionated samples, and 5,575 in non-fractionated samples. Fractionated samples thus lost around 25% of their unique precursors, whilst non-fractionated samples lost around 40%. This significant difference suggests that not only does fractionation result in more unique precursors identified, but also results in individual precursors being identified in more patients on average, giving denser, richer data sets (two sample proportion test, p-value = 5.16e-184).

Looking at figure 5, the fractionated samples generally aggregate proteins using more unique precursors (83% of proteins aggregated with two or more unique precursors), whilst a major fraction of non-fractionated proteins, 42%, are aggregated from just one unique precursor. The mean number of unique precursors used for protein aggregation was 12 vs 5 for fractionated and non-fractionated samples respectively, whilst the median was 3 and 2 respectively, a significant difference (Mann-Whitney-Wilcoxon test, p-value < 2.2e-16).

**Figure 5.**
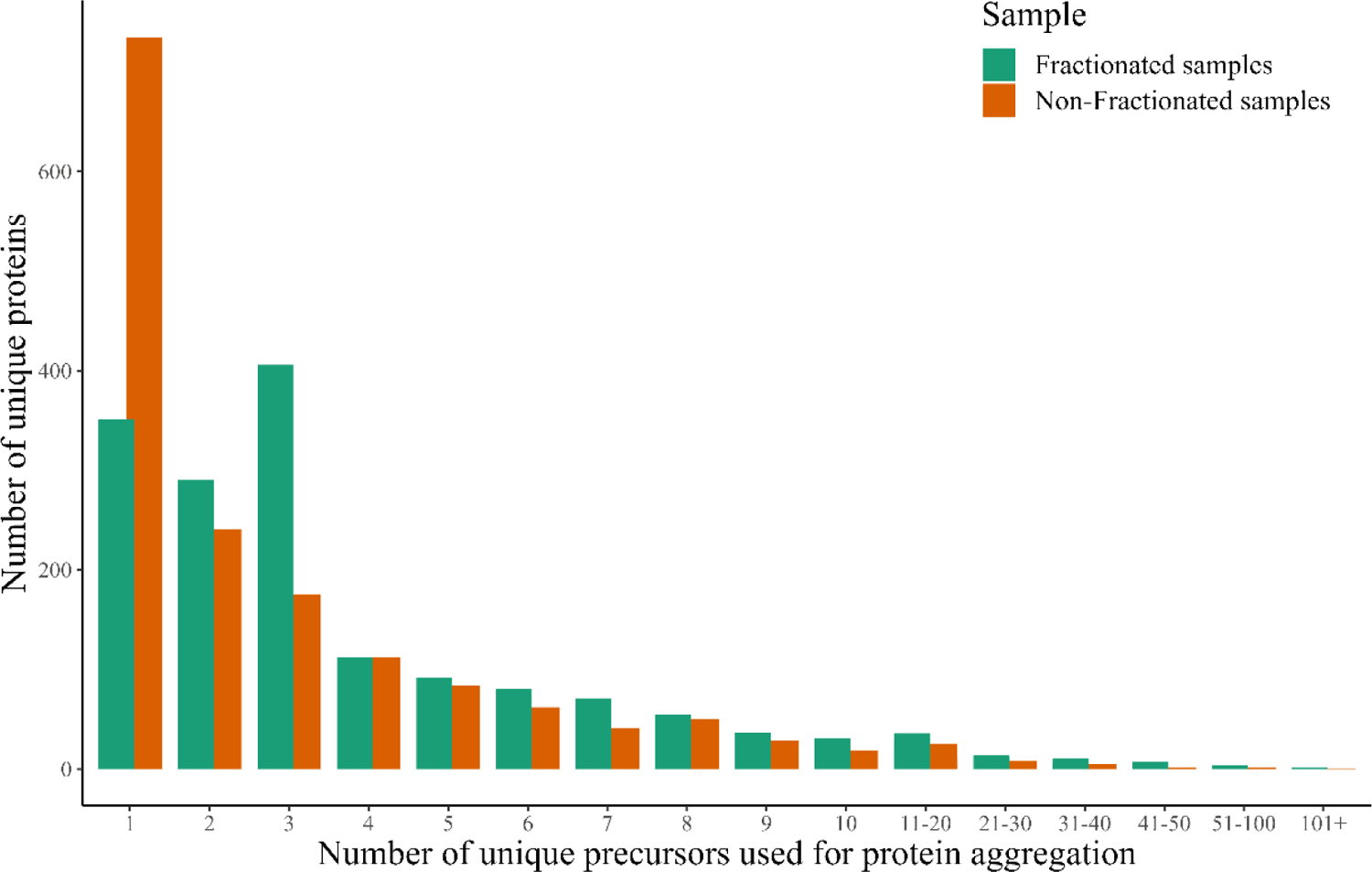
Bar plot. Bar height represents a count (y-axis) of the number of unique proteins that were aggregated with number of unique precursors as indicated on the x-axis.

Precursor abundance (calculated as MS1-based AUC) was significantly higher on average by around 32% in fractionated samples compared to non-fractionated samples (lognorm abundance 16.3 vs 15.9 respectively, two-tailed t-test, p-value <2.2e-16). Looking at figure 6A, we can see a general tendency for precursors found in fractionated samples (both uniquely and commonly identified) to be quantified at higher abundances than those in non-fractionated samples. The reduced average abundance of precursors in non-fractionated samples is not because non-fractionated samples are uniquely identifying lower abundant proteins (in fact, quite the opposite, as fractionated samples are able to uniquely identify substantially more at this lower bound), but because the lower abundant precursors found in fractionated samples are quantified with a higher abundance, potentially due to reduced competition within the MS orbitrap system.

**Figure 6.**
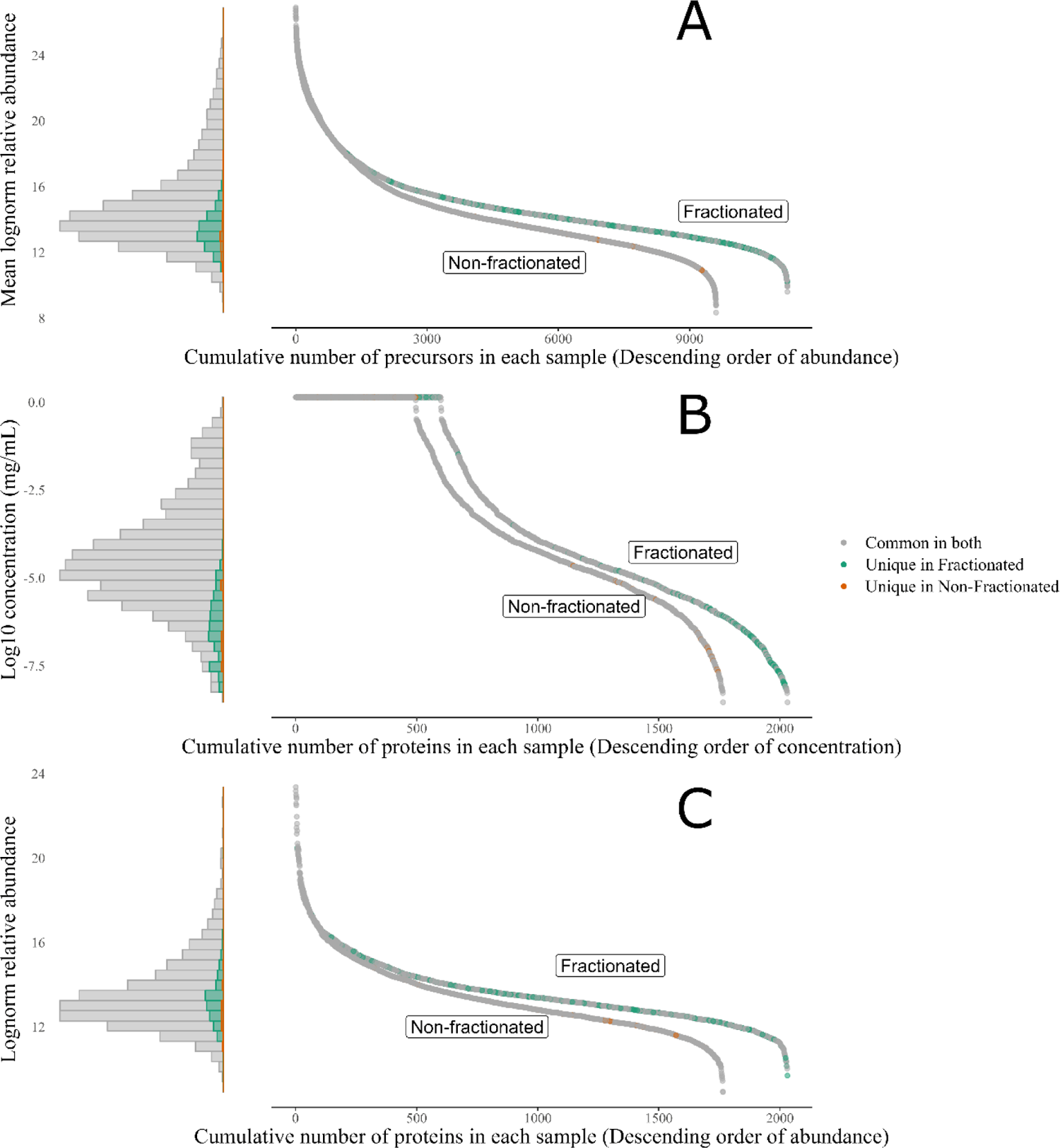
In all graphs, the left-hand side represents the data in histogram format whilst the right-hand side shows the data as a point graph, with each point representing a protein/precursor) and ranked and ordered along the x-axis by descending abundance/concentration. Proteins/precursors found in both fractionated and non-fractionated samples are marked in grey, those unique to fractionated samples are marked in green, and those unique to non-fractionated samples are marked in orange. **A**: Average abundance of uniquely identified precursors **B**: Plasmatic concentration of identified proteins in fractionated and non-fractionated samples. Proteins with a concentration of 0 mg/mL did not have a concentration listed in the Plasma Proteome Database ^35,36^, but were included to better represent total numbers of proteins in each sample. **C**: Average abundance of uniquely identified proteins.

This trend is repeated in figure 6B, which shows the plasmatic concentrations of these proteins as reported in the literature from healthy individuals (Plasma Proteome Database) ^35,36^. Again, most of the commonly identified proteins tend to be present at higher concentrations than the uniquely identified proteins, which are all almost always identified uniquely in the fractionated samples and present at lower concentrations. Fractionation of samples results in a slight enhanced detection of proteins found at lower biological concentrations (log10 concentration (mg/mL) of −4.7 vs −4.5 for fractionated and non-fractionated samples respectively, left-tailed t-test, p-value = 0.001). However, this difference is much more pronounced when filtering is applied (only precursors found in >25% of samples), here we see an average concentration of −4.4 vs −3.9 for fractionated and non-fractionated samples respectively (left-tailed t-test, p-value = 5.02e-12). Although both methodologies are able to detect similarly low abundant proteins, fractionated samples are able to detect these proteins more commonly across a cohort, with denser data allowing for far superior downstream data analysis.

Looking to the average experimental abundance of these proteins (figure 6C), we also see a slight, but significant difference in protein abundances, with fractionated proteins appearing to detect proteins at a slightly lower lognorm abundance compared to non-fractionated proteins (13.85 vs 14.03 respectively, two-sided t-test, p-value = 0.040). Inversely, when investigating common proteins (those found in both workflows), fractionated samples detect these proteins at a significantly higher abundance than in non-fractionated samples, with a mean lognorm abundance 0.31 higher in fractionated samples compared to non-fractionated samples (13.62 vs 13.31 respectively, two-tailed, paired, t-test, p-value < 2.2e-16).

Fractionated samples were collected in 7 fractions before being pooled into 3 fractions. In order to assess the potential loss of information from the pooling process. The original 7 fractions from one sample were compared to the pooled fractions and non-fractionated aliquots. Figure 7 shows the number of precursors identified in each fraction for each workflow, with the non-fractionated sample containing the highest number of precursors per aliquot, and the 7 fractions containing the least number of precursors per aliquot. Additionally, with each additional aliquot, we can observe the amount of new information introduced, in the form of unique precursors, is generally reduced, as most of the subsequent aliquots contain less and less unique precursors and more and more previously identified precursors. When looking at all aliquots of each sample, non-fractionated samples contain 6,099 unique precursors, pooled samples contain 7,165 unique precursors (13,973 total precursors across all fractions), and the original 7 fractions contain 8,477 unique precursors (24,550 total precursors across all fractions).

**Figure 7.**
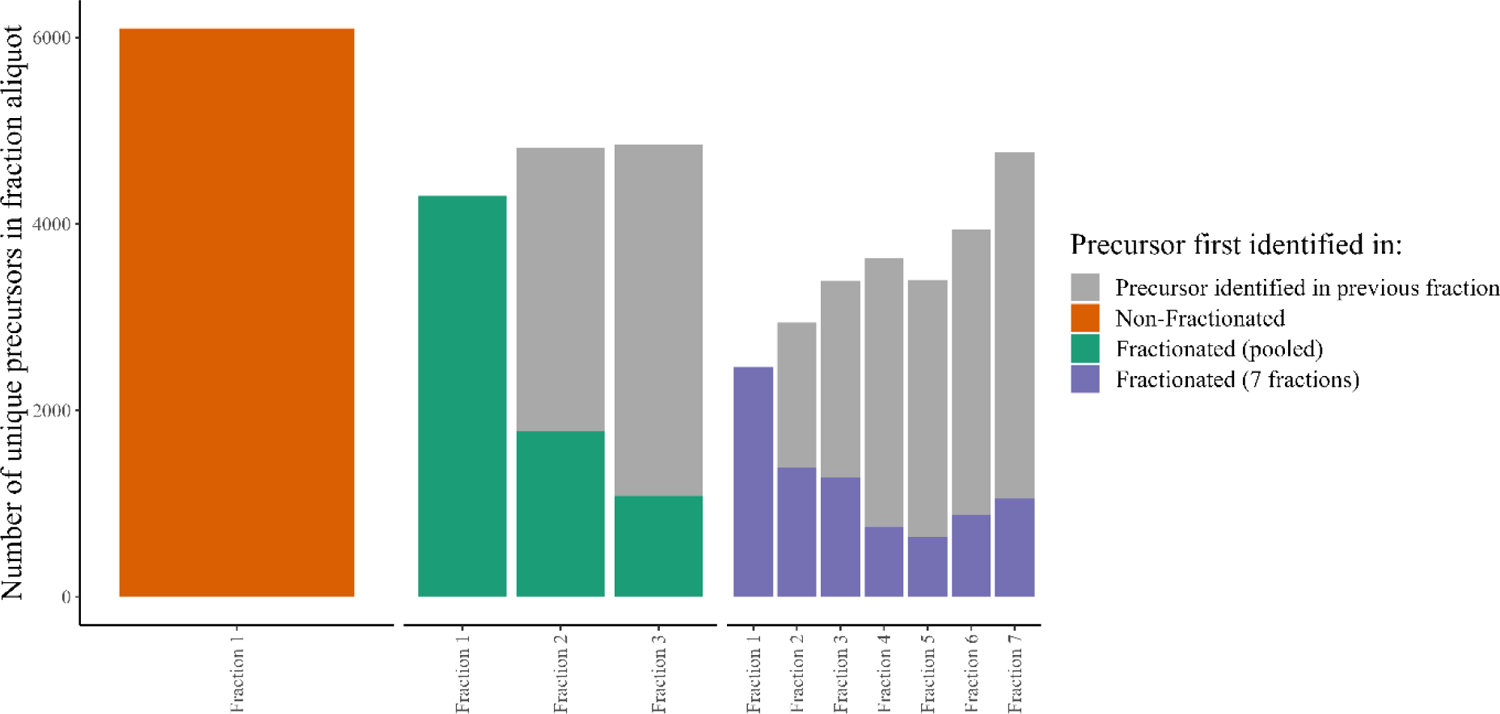
Uniquely identified precursors identified per aliquot of each sample, separated by sample type (non-fractionated, pooled fractions, 7 (non-pooled) fractions). Bar height represented the total number of unique precursors identified, whilst colour represents what proportion of those precursors are newly identified or previously identified (within each sample type). Precursors identified in previous fractions are identified in grey, new precursors are identified in either orange, green, or purple depending on sample type.

As far as how these additional precursors relate to protein identification, figure 8 shows data from the same samples processed with 3 different fractionation methods and acquired in the LC-MS/MS system in either DDA or DIA modes, in order to make a comparison between both fractionation methods as well as acquisition methods. Figure 8 shows a large increase in protein identification in fractionated samples compared to non-fractionated samples. This increased protein identification tends to be predominantly for lower abundant proteins. In terms of how the pooling process affects protein identification, we can see a 16% drop (232 proteins) from the seven fractions to the three pooled fractions. For DDA data, this drop is more pronounced, with a 24% drop in protein identification when pooling fractions (121 proteins). As expected, these proteins tend to be towards the lower end of the concentration range for each method, although both fractionation workflows are able to identify proteins in equally low concentrations.

**Figure 8.**
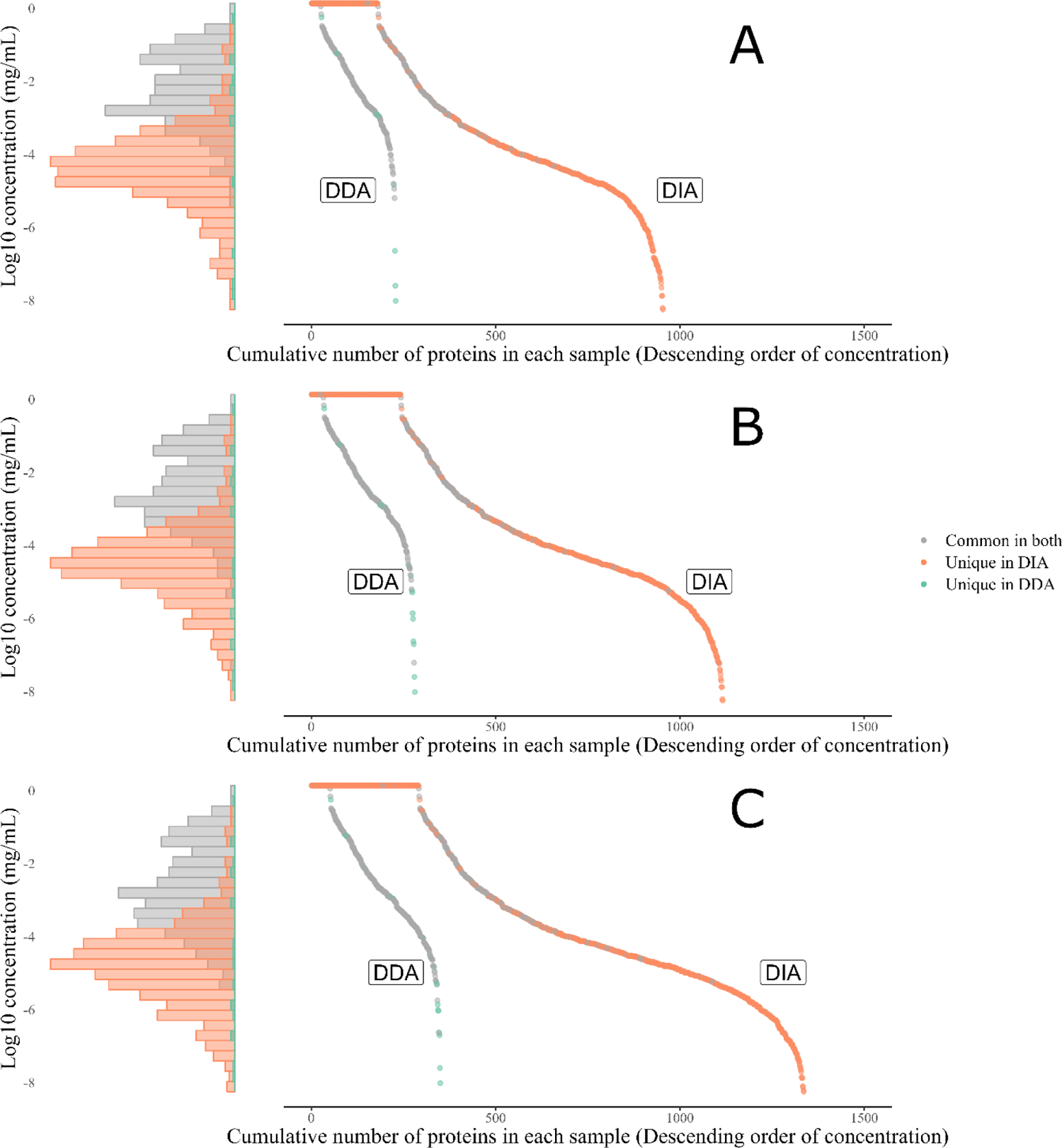
The left-hand side represents data as a histogram, whilst the left-hand side is a point graph in which each point represents a unique protein, as identified in either DDA or DIA workflows, arranged along the x-axis in descending order of plasmatic concentration (y-axis). **A** = non-fractionated, **B** = pooled fractions, and C **=** 7 (non-pooled) fractions. Proteins common to both DDA and DIA are identified in grey. Proteins unique to DIA workflows are identified in orange, and proteins unique to DDA workflows are identified in green. Proteins with a concentration of 0 mg/mL did not have a concentration listed in the Plasma Proteome Database ^35,36^, but were included to better represent total numbers of proteins in each sample.

## DDA vs DIA

Figure 8 shows a significant increase in the number of identified proteins using DIA as compared to DDA. Using DIA boosts the number of unique, detected proteins 249% for non-fractionated samples (+744 proteins), 211% for pooled fractions (+826 proteins), and 183% for un-pooled fractions (+937 proteins). Additionally, the proteins uniquely identified in samples ran in DIA mode all tend to be low-abundant proteins, and although there is some overlap between the abundances of proteins identified in DDA and DIA modes, DIA mode has a much greater abundance range of acquisition. 90% of the identified DDA proteins have an abundance between 89µg/ml and 0.058µg/ml, where as 90% of DIA proteins have an abundance between 13µg/ml and 0.0014µg/ml. Mean concentration was 30.2µg/ml and 0.996µg/ml for DDA and DIA proteins respectively.

Using the entire 20 patient cohort, figure 9 shows a direct comparison between DDA and DIA analytical modes for non-fractionated samples. More than triple the number of unique proteins are found in DIA samples as compared to DDA samples, 1,765 vs 563 respectively. Further, DIA is also seen to have slightly better coverage in terms of identification of each protein within samples, with each protein appearing in an average of 9.49 samples vs an average of 8.39 samples for DDA proteins (out of 20 samples total) (right-tail t-test, p-value = 0.0015). When considering proteins found in both DIA and DDA, the difference is further exacerbated, with these proteins being found in 17.1 DIA samples vs 9.77 DDA samples (right-tail t-test, p-value < 2.2e-16). Finally, in terms of core proteins (those proteins found within all 20 samples), DIA has more than double the number of core proteins at 303, as compared to 116 core proteins found with DDA.

**Figure 9.**
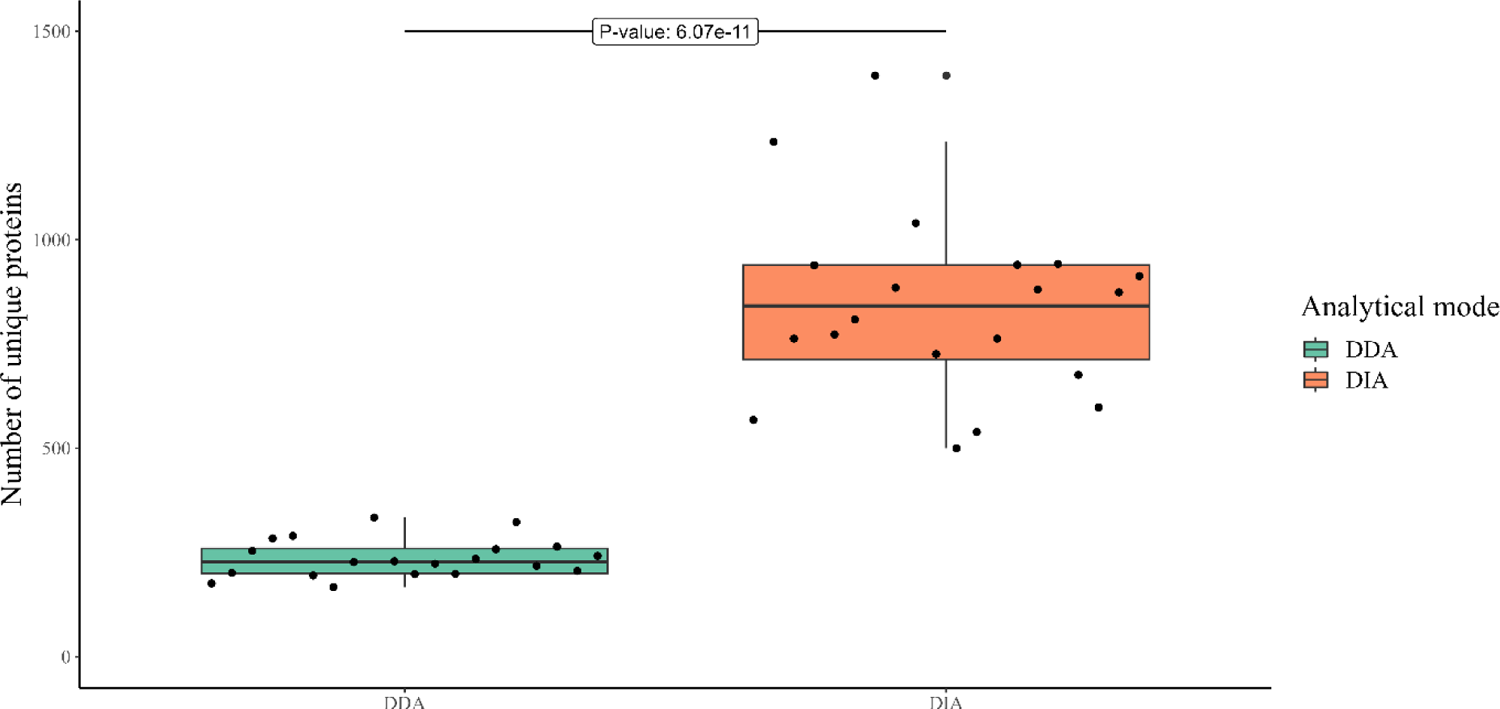
Boxplot of the number of unique proteins detected per patient in DDA and DIA modes using identical, non-fractionated samples. Points represent individual patients. Right-tailed, paired, t-test, t = 12.55, p-value = 6.07e-11

Taken together, these data would suggest that not only does DIA allow for a great increase in the number of proteins detected per sample and at lower abundances, but that it also allows for a much greater identification rate of those proteins between samples.

### Preliminary cohort analysis

As the cohort used here was part of the larger HYGIEIA cohort for a study into COVID-19, we provide here some preliminary biostatistical analyses to investigate how the fractionated samples compare to non-fractionated samples when investigating biological differences. Figure 10 shows that both methodologies can separate healthy controls and moderate COVID-19 from the respiratory failure group and severe COVID-19 which required hospitalisation, when analysed by Principal component analysis (PCA, principal components 1 and 2). The non-fractionated samples separate these four groups along PC1 and PC2, whilst the fractionated samples separate the four groups along PC1 only. Including PC3 for fractionated samples can help to further separate overlapping healthy controls and moderate COVID-19 patients to some extent. Non-fractionated samples are able to capture around 65% of the data variation in PC1 and PC2, whilst fractionated samples are only able to capture around 40% in these two components.

**Figure 10.**
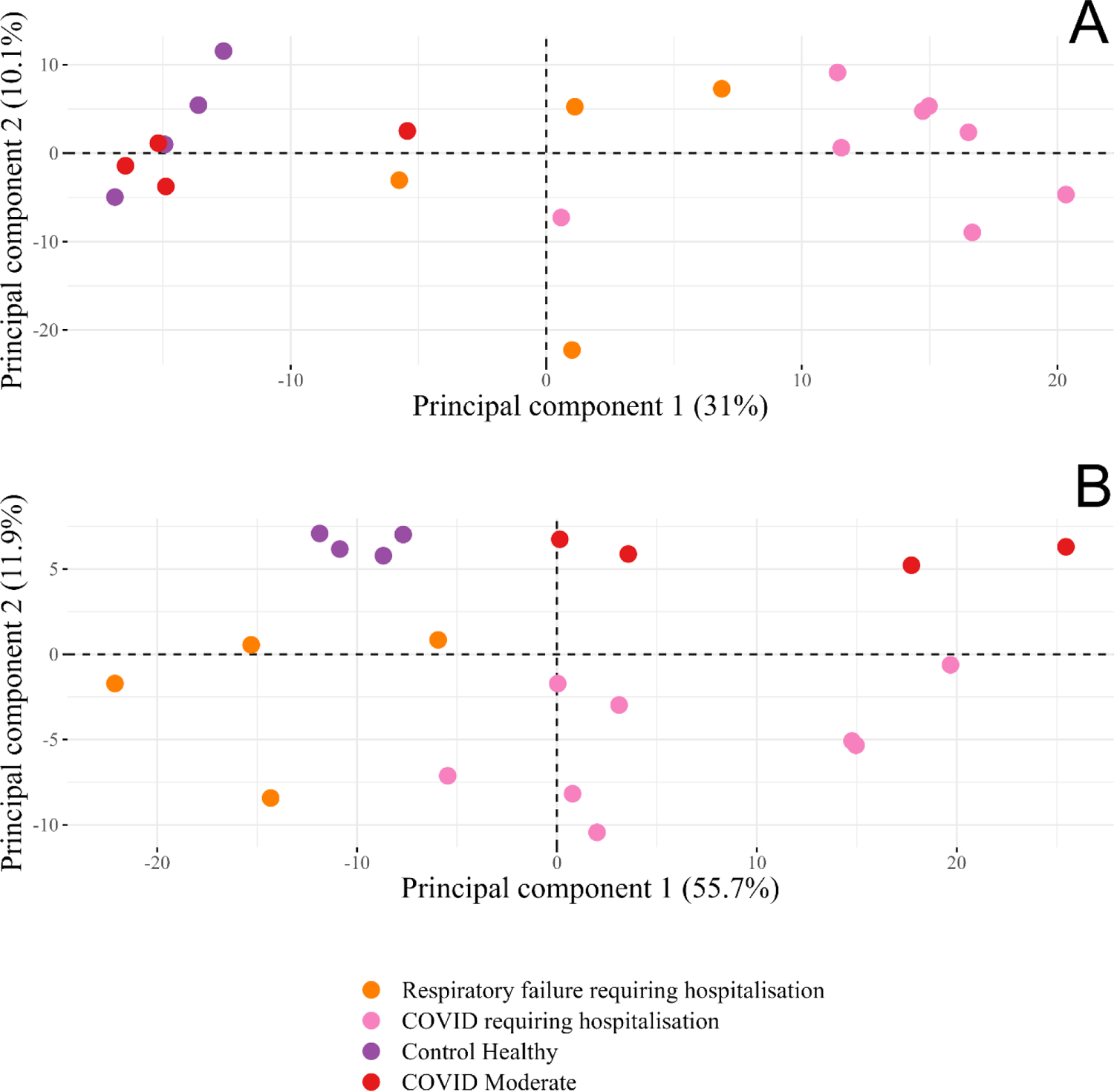
PCA of fractionated samples (A) and non-fractionated samples (B). Patients are grouped into four categories: two controls, a healthy control (purple) and a severe respiratory failure control (orange), and two COVID-19 groups, moderate COVID-19 (red) which did not require hospitalisation, and severe/critical COVID-19 (pink) which did require hospitalisation.

When looking at a differential abundance analysis (moderate COVID-19 patients vs hospitalised (severe and critical) COVID-19 patients), we see the fractionated samples (figure 11A and 10B) significantly outperform the non-fractionated samples (figure 11C and 10D) at identifying differentially abundant proteins, 264 vs 23 (two sample proportion test, p-value = 4.73e-35). Of these differentially abundant proteins, the number of COVID-19 biomarkers is also greater in fractionated samples (9 vs 5), although not at a significantly different proportion (two sample proportion test, p-value = 0.086). Finally, we are also able to detect a significantly higher number of COVID-19 associated proteins in fractionated samples (80) than in non-fractionated samples (13) (two sample proportion test, p-value = 1.67e-10). Some of the most significantly expressed proteins in fractionated samples were tyrosine 3-monooxygenase (YWHAE), reticulon 4 (RTN4), pleckstrin (PLEK), profilin 1 (PFN1), and haptoglobin (HP).

**Figure 11.**
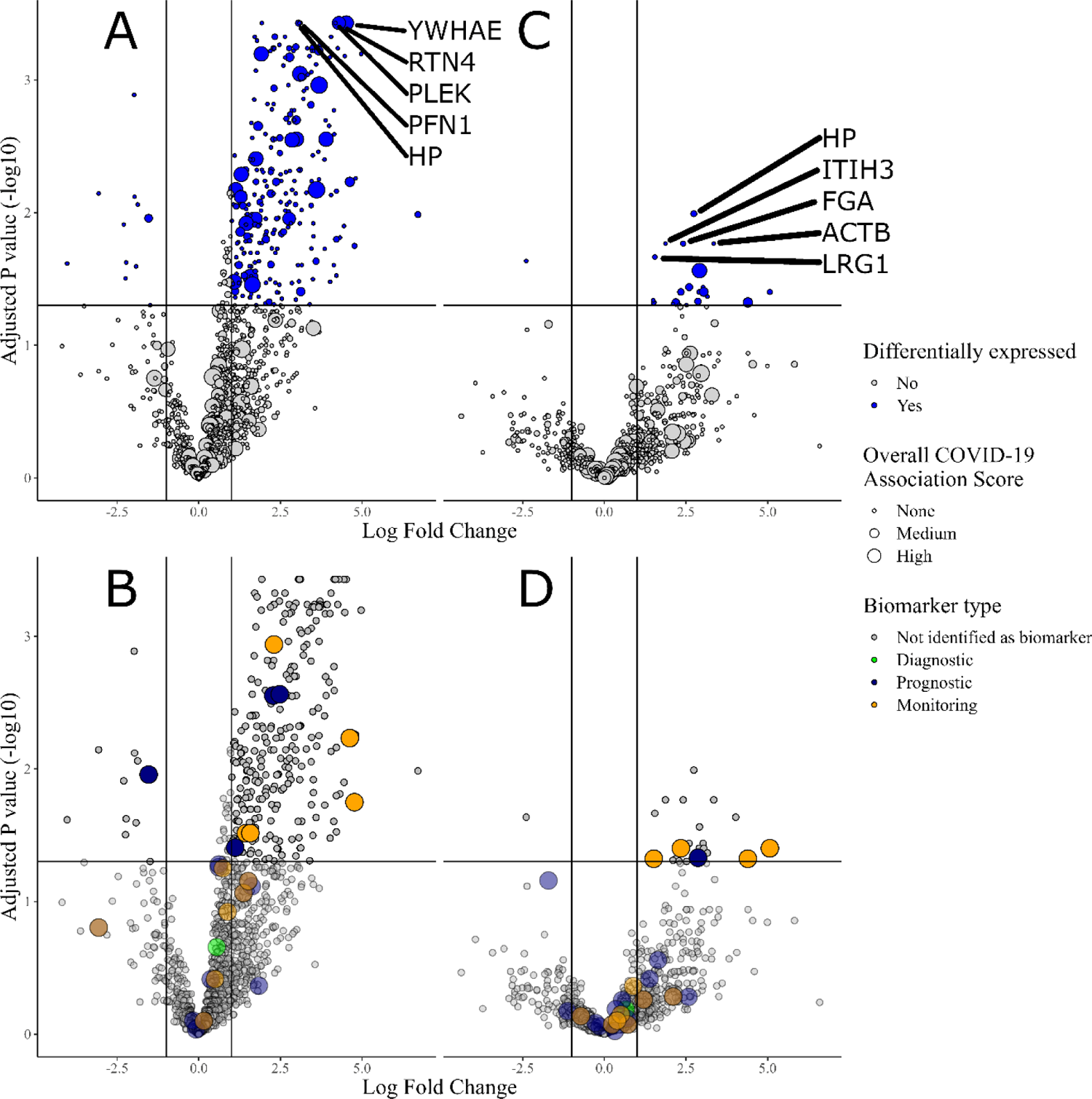
Volcano plot of differentially expressed proteins between non-hospitalised COVID-19 patients and hospitalised COVID-19 patients. **A** and **B** shows differentially expressed proteins in fractionated samples, whilst **C** and **D** shows differentially expressed proteins in non-fractionated samples. **A** and **C** show differentially expressed proteins (absolute log fold change > 1 and adjusted p-value <= 0.05) coloured in blue, and point size represents overall COVID-19 association score as calculated via OpenTargets platform ^38^. **B** and **D** highlight points which have been identified as either diagnostic, prognostic, or monitoring biomarkers (point size is irrelevant and is used to better highlight the biomarkers). OncoMX COVID-19 biomarker database used for biomarker identification ^37^.

## Discussion

Our data unequivocally shows fractionation of samples leads to superior results when looking at precursors and proteins identified, reliability of detection of these precursors and proteins across samples, and downstream differential analysis between groups. The benefits of plasma prefractionation methods have long been shown using DDA methodologies, and extensive fractionation techniques were often employed in deep plasma proteome studies to overcome the limitations of DDA techniques. In our data, we show an average of around 236 proteins identified per patient with DDA (non-fractionated), which is roughly in line with what has been reported in other studies ^39^. In contrast, our DIA samples were able to detect an average of 838 unique proteins per patient, and up to almost 1,500 in some patients (figure 9). On top of this, we can see clear improvements in terms of numbers of core proteins, and reliability of protein identification between samples, as highlighted in figure 3.

Figure 8 allows for direct comparison of results for samples analysed using a methodology that incorporates no fractionation, pooled fractions, or no fraction pooling. Regardless of fractionation methodology, DIA identifies and quantifies significantly more proteins than DDA, with an average improvement of over 200% additional unique proteins.

This improvement in performance can be explained through a number of reasons. To begin, the data collected during DIA are much more detailed and comprehensive to that collected during DDA (figure 1). Traditional DDA methods by their nature exclude a large number of precursors from being selected for further MS2 analysis. Instruments select for the top N most abundant precursor ions for fragmentation during each scan, followed by a period of dynamic exclusion, preventing fragmentation of multiple precursors and thus analysing a sub-set of the data in order to reduce data complexity ^40^. Such an approach is stochastic, as small differences at any point of the run may impact precursor selection and exclusion, with these changes proliferating other changes further down the run. In contrast, DIA selects precursors for fragmentation independently of the MS1 precursor scan, as the mass spectrometer is programmed to select, group, and fragment precursors across multiple, predefined m/z windows (usually 5-25 m/z). This prevents small changes having snowballing effects on data acquisition, improving reproducibility and reliability, as well as allowing for the fragmentation of both low and high abundant precursors.

Reducing data complexity using DDA was previously critical due to limited bioinformatic tools which were unable to handle co-fragmentation and assumed the product ions seen in MS2 spectrum are derived from a single, isolated precursor. These spectra were then matched to the appropriate precursors using *in-silico* predicted peptides generated from FASTA files ^41–43^. In comparison to this, for precursor-centric methods such as DIA-NN, each precursor is detected by querying a precursor in the library and searching the MS data for suitably matching spectra. In this way they excel at interpreting the complex spectra found in DIA data due to co-fragmentation being explicitly tolerated as single spectrum can be evidence of detection for multiple precursors. Logically, this gives DIA much higher data density, allowing it to identify more precursors for a given number of spectra than DDA, which is only able to identify one precursor per spectrum.

The use of fractionation in the present study, increases the average number of identified proteins to 1,321, and just over 1,700 proteins for certain patients. In addition, we see better inter-patient data uniformity in fractionated samples as the number of core proteins, proteins detected in all patients, increases by 88%, from 303 to 570. Practically, the proteins identified in fractionated samples include almost all proteins identified by non-fractionated samples as well (only 30 out of 2,061 proteins were unique to non-fractionated samples), indicating there is minimal gain in running non-fractionated samples alongside fractionated samples for the purpose of improving detection, thus allowing for a reduced number of injections to 3 per subject.

Of the additional proteins identified in fractionated samples, most of these were towards the lower end of the abundance range. There are two main reasons for this, firstly fractionation strategies result in less complex samples with improved peptide separation, and secondly, splitting a sample into multiple fractions that are then injected separately increases the total amount of sample being run. This increase in protein identification at lower abundances is due to increased sensitivity of precursor identification. Fractionation of the sample allows for less interference of competing peptides for fragmentation and produces clearer and more sensitive MS2 spectra for bioinformatic analysis. Not only does this lead to identification of lower abundant proteins, but also gives rise to the identification of multiple precursors derived from singular proteins. This can be seen in figure 5, whereby proteins identified in fractionated samples are aggregated less from single unique precursors, and more from multiple unique precursors. This allows for a certain level of data redundancy, whereby missing a single precursor in a run will not necessarily mean a protein is not identified, which contributes to both an increase in the number of core proteins and a more representative quantitation.

In comparison to more traditional DDA proteomic studies utilising fractionation, sodium dodecyl-sulfate (SDS) and C18 peptide fractionation approaches can identify between 3,254 and 4,219 proteins by analysis of either 24 gel bands for the SDS approach, or 84 fractions concatenated into 24 fractions for the C18 approach^44^. One of the most in-depth studies into the plasma proteome was able to identify an average of 4,641 proteins per patient using a workflow consisting of a two-step depletion of the top 50-100 most abundant proteins, followed by reverse phase peptide fractionation totalling 86 fractions pooled into 30 fractions ^45^. Other examples of such studies have employed fractions sizes of 72 ^46^, 60 pooled into 18 ^47^, 6 ^48^, or 18 ^49^, and have identified a total of 1,917, 1,069, 567, and 169 proteins across all patients/samples.

However, the methodologies for these studies, and other studies utilising extensive fractionation, are complex, expensive, time-consuming, and unlikely to be practical for many clinical applications and settings, which would thus restrict their utility outside of well-funded research environments. In terms of MS run times, the above studies range between 468 and 3,600 minutes per patients/sample. In contrast, we have presented above a stream-lined, cost efficient protocol which can be deployed for both exploratory research and clinical applications. This protocol includes easily sourced reagents and kits, shorter experimental time compared to other such studies, and by producing only 7 fractions concatenated into 3 pooled fractions only, an MS runtime of just 360 minutes per sample for an average of over 1300 proteins identified per patient. Reducing the cost and time investment of proteomic experiments gives lower funded projects and laboratories the opportunity to introduce large scale MS proteomics into their experiments, as well as allowing current MS proteomic labs the ability to increase their sample and cohort sizes.

Pooling 7 fractions into 3 fractions reduced unique precursor identification and protein identification by around 15%, but improved the efficiency of each fraction in terms of total and unique precursors per fraction (figure 7). The dropped precursors and proteins were from the lower end of the abundance range, although both techniques were able to detect proteins/precursors at equally low abundances. Total MS run time by pooling fractions was reduced by almost 60%, from 840 minutes to 360 minutes. Running un-pooled fractions does result in superior protein identification, yet depending on the goals, funding, and time constraints of the project, the slight drop in capability may be worth it in exchange for the massive reduction in running time and costs, especially as extensive fractionation results in diminishing returns in terms of unique precursors per subsequent fraction.

Previously, such extensive fractionation was required in DDA experiments due to its higher missing values, lower reproducibility, lower quantitative accuracy, and lower protein identification ^22,50,51^. Now however, the use of DIA can be used to either greatly improve results when paired with the same protocol, or, the work presented here demonstrates that simpler, faster, and more cost-effective protocols, when used in conjunction with DIA proteomic methods, can yield similarly large and deep proteome coverage as more complex DDA proteomic approaches. However, the work here also demonstrates limits to the how simple the protocol may become, even with the vast improvements offered by a DIA approach, the use of fractionation is a must for studies looking to investigate differentially expressed proteins, which may act as pathophysiological signals or biomarkers.

For the COVID-19 preliminary cohort used here, figure 10 shows that although non-fractionated samples are able to find variations between groups that can be used to separate them, figure 11 shows that further investigations via differential expression analysis yield sub-par results as compared to fractionated samples.

In both sample types, figure 10 shows that each are able to separate severe respiratory failure, both COVID-19 derived and non-COVID-19 derived, from moderate or non-infection states. In terms of differentially expressed proteins between moderate and severe/critical COVID-19, both fractionated and non-fractionated samples were able to identify 264 and 23 proteins respectively. Of these, CRP, SAA1, CST3, VWF and S100A9 were found to be significantly upregulated in both samples. These proteins have all been identified as potential biomarkers within other studies; increases have been associated with higher disease severity, more severe complications, and in the progression of patients that require mechanical ventilation ^52–56^. In contrast, albumin (ALB) was found to be significantly downregulated in fractionated samples, which is line with other studies that have identified hypoalbuminemia as an independent prognostic biomarker that has been significantly associated with severe and critical disease conditions and may predict fatality risk in critically ill COVID-19 patients ^57,58^. In addition to this, fractionated samples also identify significantly increased expression in hospitalised COVID-19 patients (as compared to non-hospitalised patients) of previously identified biomarkers CSF1, SELP, and LGALS3 ^53,56,59,60^. For fractionated samples, four biomarkers, SFTPD, GOT1, MB, and PF4 had a positive logfold change over 1, while KRT19 had a negative logfold change over 1, but all lacked significance, perhaps due to the small cohort size in the present study. Upregulation of SFTPD, GOT1, and MB was previously found to be associated with severe disease and follow the same trend in this study ^53,61–63^.

In fractionated samples, the most significant proteins were YWHAE, RTN4, PLEK, PFN1, and HP; of these, HP and PFN1 were also found to be differentially expressed in non-fractionated samples, albeit at lower significances. Two biomarkers (HP and YWHAE) have been associated with COVID-19 through the OpenTargets aggregated association score utilising various data sources (Europe PMC, Expression Atlas, IMPC, PROGENy, etc) ^38^. PLEK, which was not associated with COVID-19 using the above-mentioned scoring system, is involved in G protein-coupled receptor signalling pathway, actin cytoskeleton organisation, and positive regulation of supramolecular fiber organization. Consistent with this study, PLEK was also found to be significantly differentially expressed in a study investigating SARS-CoV-2 infection and negative controls ^64^. Additionally, an investigation into IFN signalling in the nasopharynx of SARS-CoV-2-infected individuals found that PLEX was significantly upregulated as part of the IFN-mediated antiviral response ^65^.

Focusing on non-fractionated samples for a moment, ST3GAL6 was the only differentially expressed protein that was not differentially expressed in fractionated samples (Non-fractionated: logFC –2.39, adjusted p-value 0.023. Fractionated: logFC −0.23, adjusted p-value 0.73.). ST3GAL6, a member of the sialyltransferase family, is not a known biomarker for COVID-19 and has not yet been associated with COVID-19 per the OpenTargets Platform or in the general literature.

Some of the lowest abundant proteins that were shown to be differentially expressed in fractionated samples include, GALNT12, TTC36, and CLEC11A, usually present at concentrations of 15ng/L, 20ng/L, and 45ng/L respectively. For non-fractionated samples, ST3GAL6, MSN, YWHAZ were the top 3 lowest abundant proteins at 3µg/L, 340µg/L and 760µg/L respectively. The median concentration of differentially expressed proteins (as reported in the Plasma Proteome Database) for fractionated samples was around 1/50^th^ the median concentration of differentially expressed proteins for non-fractionated samples, at 85µg/L vs 4.3mg/L respectively ^35,36^. Fractionation leads to a higher number of differentially expressed proteins, which are found at a much lower average biological concentration. Fractionation, due to its ability to enhance the detection of proteins across a cohort, appears indispensable in identifying low abundant proteins crucial for comprehensive proteomic analysis.

Among those low abundant proteins identified in fractionated samples, are some that have been associated with a role in COVID-19 disease, such as FGFR4 (4.3µg/L), which acts as a cell-surface receptor for fibroblast growth factors and is one of the targets of NINTEDANIB, a small molecular drug currently undergoing phase III trials for treatment of COVID-19 via inhibition of mesenchymal cell activation and virus-induced pulmonary fibrosis (National Library of Medicine [NLM], NCT04541680). Another such protein MGAT1 (7.2µg/L), is involved in the maturation pathway of the SARS-CoV-2 spike (S) protein, though the glycosylation of the host-derived glycans which cover the proteins surface ^67^.

## Conclusions

In conclusion, we have presented an exploratory proteomic protocol with a simplified fractionation procedure and boosted by the utilisation of DIA methodologies. This pilot study illustrates the potential clinical applicability of this protocol when used in conjunction with a DIA bioinformatics pipeline. Using this protocol, hundreds of extremely low abundant proteins usually present at concentrations as low as 15ng/L have been identified as significantly differentially expressed. In addition to this, we show how the use of fractionation methods in addition to DIA technologies can significantly boost unique protein identification within samples and greatly improve identification of the same proteins between samples. The method discussed in this study was able to identify an average of 1,321 proteins per patient using plasma samples collected as part of a 20-patient COVID-19 cohort. Across the entire cohort, 2,031 total unique proteins were identified. The protocol employs the use of readily-available kits and materials for protein depletion, digestion, and fractionation for a time-efficient and simple sample preparation. 7 fractions are then concatenated into 3 pooled fractions, for a total MS run time of 360 minutes. Utilising such a protocol can significantly reduce the cost, time, and complexity required in performing exploratory proteomic studies on human plasma and is currently being applied in our setting to a large, 200 +, multi-timepoint patient cohort in a multi-omics context ^29^.

## Author Contributions

The manuscript was written through contributions of all authors. All authors have given approval to the final version of the manuscript. Conceptualization, J.C.Y., J.-L.B., L.G., J.d.G., L.B. and L.E.; methodology, B.W., S.P.d.R., M.L., D.V., and L.E.; investigation, B.W., J.C.Y., J.d.G., J.-L.B., P.D.C., J.-F.C., J.P.D., L.G., V.H., S.J., B.K., S.P.d.R., D.V., L.E. and L.B.; resources, J.C.Y., J.-L.B., L.G., J.d.G., V.H., B.K., J.-F.C., L.B. and L.E.; writing—original draft preparation, B.W., S.P.d.R, and L.E.; writing—review and editing, J.C.Y., J.d.G., J.-L.B., P.D.C., J.-F.C., J.P.D., L.G., V.H., S.J., B.K., L.B., and D.V.; supervision, J.C.Y., J.-L.B., L.G., J.d.G., L.B. and L.E.; project administration, J.-L.B. and J.C.Y.; funding acquisition, J.C.Y., J.-L.B., L.G., J.d.G., L.B. and L.E. All authors have read and agreed to the published version of the manuscript.

## Notes

Informed consent was obtained from all subjects involved in the study. The study is conducted in accordance with the Declaration of Helsinki, and approved by the Institutional Review Board (or Ethics Committee) of Cliniques Universitaires Saint Luc (comité éthique hospital-facultaire) (protocol code 2021/30DEC/543, date of approval: 30 December 2021).

## Supporting Information

Appendix 1: R script detailing data processing, figure generation, and statistical analysis (R File).

## ACKNOWLEDGMENT

The investigators extend their heartfelt thanks to all the patients actively engaged or intending to participate in this clinical study. Special appreciation is also extended to the clinical medical research coordinators at Cliniques Universitaires Saint Luc for their invaluable support in patient recruitment and the collection of biological samples.

## ABBREVIATIONS

DDA: Data-dependent acquisition

DIA: Data-independent acquisition

LC-MS/MS: Liquid chromatography tandem mass spectrometry

COVID-19: Novel coronavirus disease 2019

LC: Liquid chromatography

MS: mass spectrometry

LC-MS: Liquid chromatography mass spectrometry

RT: Room temperature

RPM: Rotations per minute

TEAB: triethylammonium bicarbonate

ACN: Acetonitrile

TFA: Trifluoroacetic acid

APD: Advanced peak detection

HPLC: High-performance liquid chromatography

FWHM: Full width at half-maximum resolution

HCD: Higher energy collisional dissociation

AGC: Automatic gain control

FDR: False discovery rate

HYGIEIA: Hypothesizing the genesis of infectious diseases and epidemics through an integrated systems biology approach

AUC: Area under the curve

PCA: Principal component analysis

PC: Principal component

QC: Quality control

SDS: Sodium dodecyl-sulfate

logFC: Log fold change

NLM: National library of medicine

## Availability of data

Data are available via ProteomeXchange with identifier PXD047901

## For Table of Contents Only

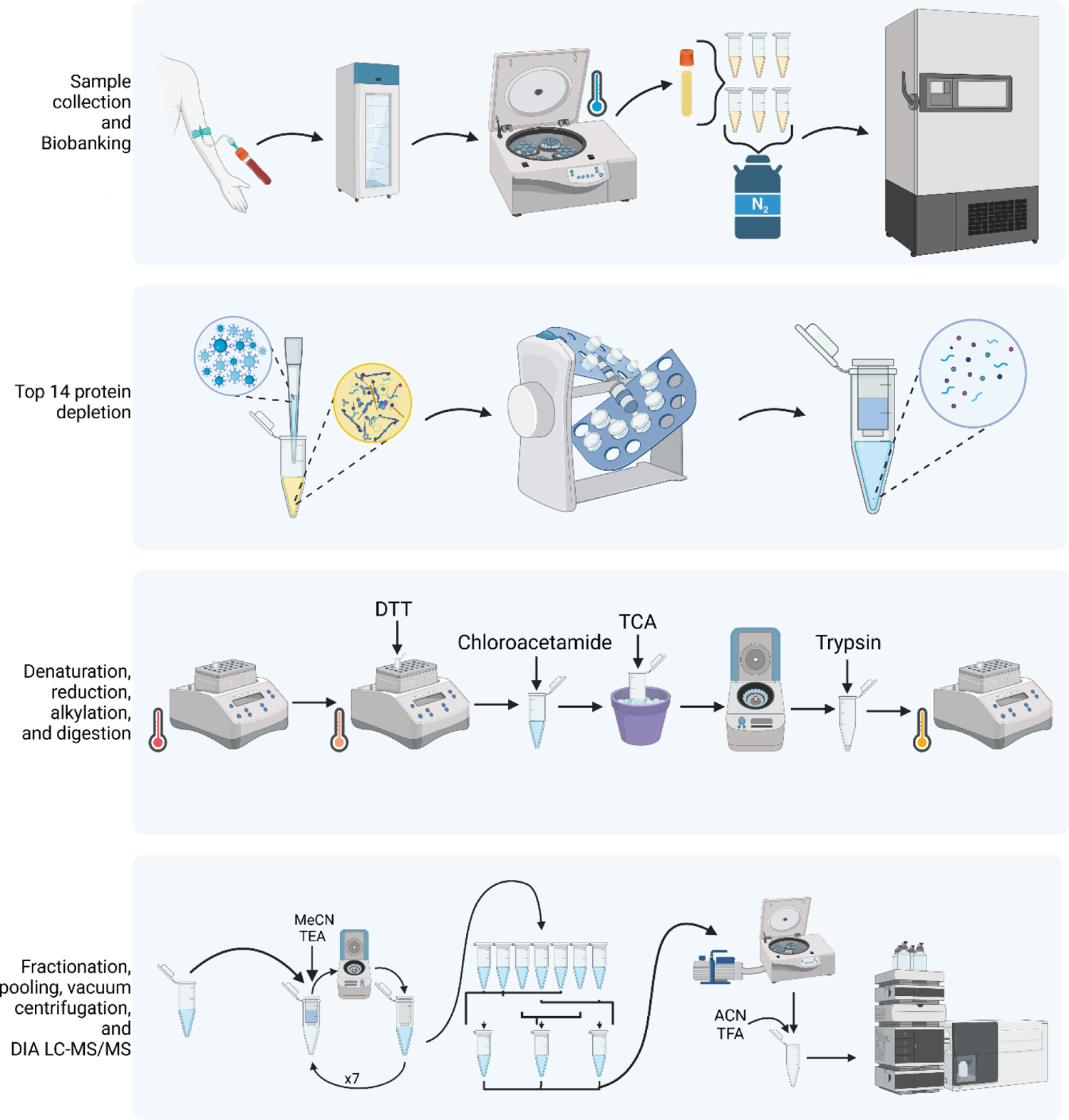

